# A Spectrum of Free Energy Landscape Topologies Encodes Chromatin Polymorphism and Phase Separation

**DOI:** 10.64898/2026.06.19.733383

**Authors:** Yifang Chen, Jan Huertas, M. Julia Maristany, Kieran Russell, Mingyuan Zhang, Stephen E. Farr, Jorge R. Espinosa, Rosana Collepardo-Guevara

## Abstract

Chromatin exhibits structural polymorphism, yet how such substantial structural heterogeneity is encoded in its underlying free-energy landscape remains unknown. Here we map the folding landscapes of tetra-nucleosome arrays using adaptive Markov state models constructed from near-atomistic coarse-grained simulations. We find that chromatin is not governed by a single folding funnel analogous to those of many globular proteins. Instead, it spans a continuum of free-energy landscape topologies, ranging from funnelled to rugged to flat. Remarkably, phase separation reshapes chromatin’s free-energy landscape by stabilising conformations that maximise intermolecular connectivity, demonstrating that the landscape is determined not only by intrinsic molecular parameters but also by the environment and intermolecular interactions. Linker DNA geometry selects among landscape regimes through geometric frustration of zig-zag nucleosome stacking. At short and intermediate linker lengths, canonical 10*N* bp linkers produce funnelled landscapes dominated by compact zig-zag structures, whereas 10*N* +5 bp linkers generate rugged landscapes with multiple competing minima. Long linkers relieve frustration and flatten the landscape into a broad entropic basin. This control operates with single-base-pair precision, such that changing linker length by one base pair is sufficient to reorganise both the thermodynamics and kinetics of chromatin folding. DNA sequence provides a second level of control by modulating linker deformability, but only when zig-zag geometric frustration is present. These results establish free-energy landscapes as a framework for understanding the physical origin of chromatin polymorphism and its coupling to phase separation.

## INTRODUCTION

The eukaryotic genome is organised hierarchically, from nucleosomes formed by DNA wrapped around histone octamers, to chromatin fibres—polymers of nucleosomes connected by linker DNA—and nuclear compartments [1**?**]. This hierarchical organisation plays a central role in regulating gene expression, DNA replication, and DNA repair. Yet chromatin is remarkably polymorphic [2–8]. Even a single nucleosome array can adopt structures ranging from compact ordered zig-zag fibres to irregular, locally disordered conformations [2, 6, 9–12]. In the physics of self-organising systems, such behaviour is naturally described by an energy landscape whose topology encodes the number, stability, and connectivity of competing structural states. For many globular proteins, folding is governed by a funnelled landscape that guides the chain toward a unique native state [13–16]. Whether chromatin is governed by a single universal landscape analogous to the folding funnels of globular proteins, or instead by multiple classes of landscape topology that encode distinct structural ensembles, remains unknown.

Chromatin’s structural diversity is both pervasive and finely tunable [2–4, 17]. Among the many parameters that regulate chromatin structure, the length and sequence of the linker DNA connecting adjacent nucleosomes are decisive, together with histone composition and extrinsic factors such as ionic strength and binding of architectural proteins including, for instance, H1, HP1, and Oct4 [9, 11, 18–21]. Linker length is particularly powerful because it sets the relative orientation of successive nucleosomes and thereby partitions their interaction preferences between intra- and inter-fibre contacts [9, 10].Canonical linkers with lengths of 10*N* bp (*N* being an integer) promote compact zig-zag folding that self-saturates the strongest face-to-face nucleosome contacts within a fibre, whereas non-canonical 10*N*+5 linkers geometrically frustrate this self-saturation, since no single fibre configuration can simultaneously satisfy all its face-to-face contacts, exposing interaction sites for intermolecular networks [9, 10, 22]. Thus, these same geometric distinctions control not only the structure of single fibres but also the propensity of chromatin to phase separate into condensates [10, 22, 23]. Yet despite extensive knowledge of how linker DNA geometry regulates chromatin compaction, nucleosome interaction patterns, and phase separation, how these behaviours are encoded in the underlying free-energy landscape—in the number, depth, and connectivity of the metastable basins that jointly determine thermodynamic stability, kinetic accessibility, and phase behaviour—remains unexplored.

Molecular dynamics (MD) simulations offer a natural route to address this gap by connecting chromatin conformations to the free-energy landscapes, metastable states, and kinetic pathways that govern their stability and interconversion. Because chromatin is a charge-rich poly-branched polymer its free energy landscape has proven difficult to be sampled exhaustively [11, 22]. Therefore, using coarse-grained models, which reduce the complexity of the system, paired with enhanced sampling techniques is essential [11, 17]. Adaptive sampling further improves coverage of conformational space by iteratively re-initialising simulations from under-sampled regions [24–29]. The simulation trajectories can then be used to construct Markov state models (MSMs), which provide a quantitative framework for estimating equilibrium populations and transition kinetics from molecular trajectories [30–35]. To construct MSMs, high-dimensional MD trajectories are often projected onto low-dimensional representations that capture the slow dynamical modes of the system. Dimensionality reduction approaches such as time-lagged independent component analysis (tICA) [36, 37] and non-linear reaction-coordinate learning methods such as flow matching for reaction coordinates (FMRC) [38] can therefore be used to generate compact kinetic spaces suitable for state discretisation.

Here, we focus on tetra-nucleosome arrays—the minimal chromatin unit that resolves both (*i*, *i* + 2) and (*i*, *i* + 3) nucleosome stacking—using simulations with the near-atomistic coarse-grained model OpenCGChromatin [22], which retains nucleotide- and residue-level detail while remaining computationally tractable. To broadly sample chromatin conformational space, we initialized extensive MD simulations from diverse structures generated by Debye length Replica Exchange MD (D-REMD). To further improve conformational coverage, we employed MSM-guided adaptive sampling, accumulating more than 920 independent simulation trajectories for each tetra-nucleosome system. These trajectories were then used to construct MSMs in FMRC reaction-coordinate space [38], enabling kinetic models that more effectively preserve slow dynamical processes. We systematically vary linker DNA length and sequence and, finally, extend the analysis from isolated fibres to chromatin condensates.

Our results reveal that chromatin is characterised by a continuum of free-energy landscape topologies—from funnelled, to rugged, to flat—selected by linker DNA geometry through geometric frustration of nucleosome stacking. This control operates with remarkable sensitivity. That is, single-base-pair changes in linker length are sufficient to reorganise the topology of the landscape and, consequently, the thermodynamic stability and kinetic accessibility of chromatin conformations, whereas DNA sequence modulates the landscape only when geometric frustration is present. Remarkably, the coupling between chromatin folding and phase separation is reciprocal. The condensed phase re-shapes the free-energy landscape by selectively stabilising partially expanded chromatin conformations that maximise intermolecular interactions, demonstrating that the landscape is determined not only by the intrinsic parameters of an individual fibre but also by its molecular environment and interaction partners. Chromatin polymorphism therefore arises not from stochastic disorder, but from a tunable free-energy landscape whose topology links base-pair-scale DNA mechanics to chromatin structure, dynamics, material properties and phase-separation behaviour.

## RESULTS

### Adaptive sampling and data-driven methods for MSM construction reveal chromatin folding dynamics

To characterise the thermodynamics and kinetics of chromatin-fibre folding, we employed an iterative, data-driven workflow that combines adaptive sampling, feature optimisation, low-dimensional reaction-coordinate learning, and MSM construction (Figure 1). The tetra-nucleosome arrays were modelled using OpenCGChromatin, a near-atomistic coarse-grained chromatin model [22] that retains residue- and nucleotide-level resolution (Figure 2A; Methods). The model, implemented in OpenMM, has been validated against experimental sedimentation velocity analyses of 12-nucleosome fibres with varying linker DNA lengths [39, 40], as well as cryo-EM [41], X-ray crystallography [42], and cryo-ET structures of chromatin arrays [43]. OpenCGChromatin accurately captures chromatin properties while exploiting GPU acceleration [44]. Using this model, we constructed tetra-nucleosome arrays spanning a wide range of linker lengths, from very short 15 bp linkers (NRL 162 bp), 20 bp linkers (NRL 167 bp), 22 bp linkers (NRL 169 bp), 25 bp linkers (NRL 172 bp), and 30 bp linkers (NRL 177 bp), up to long 58 bp linkers (NRL 205 bp). To probe linker-length dependence more finely, we also generated arrays with linker lengths from 25 to 30 bp in 1 bp increments, along with linker-sequence variants (`polyA` and `polyAT`), to capture sequence-dependent structural heterogeneity.

**FIG. 1.**
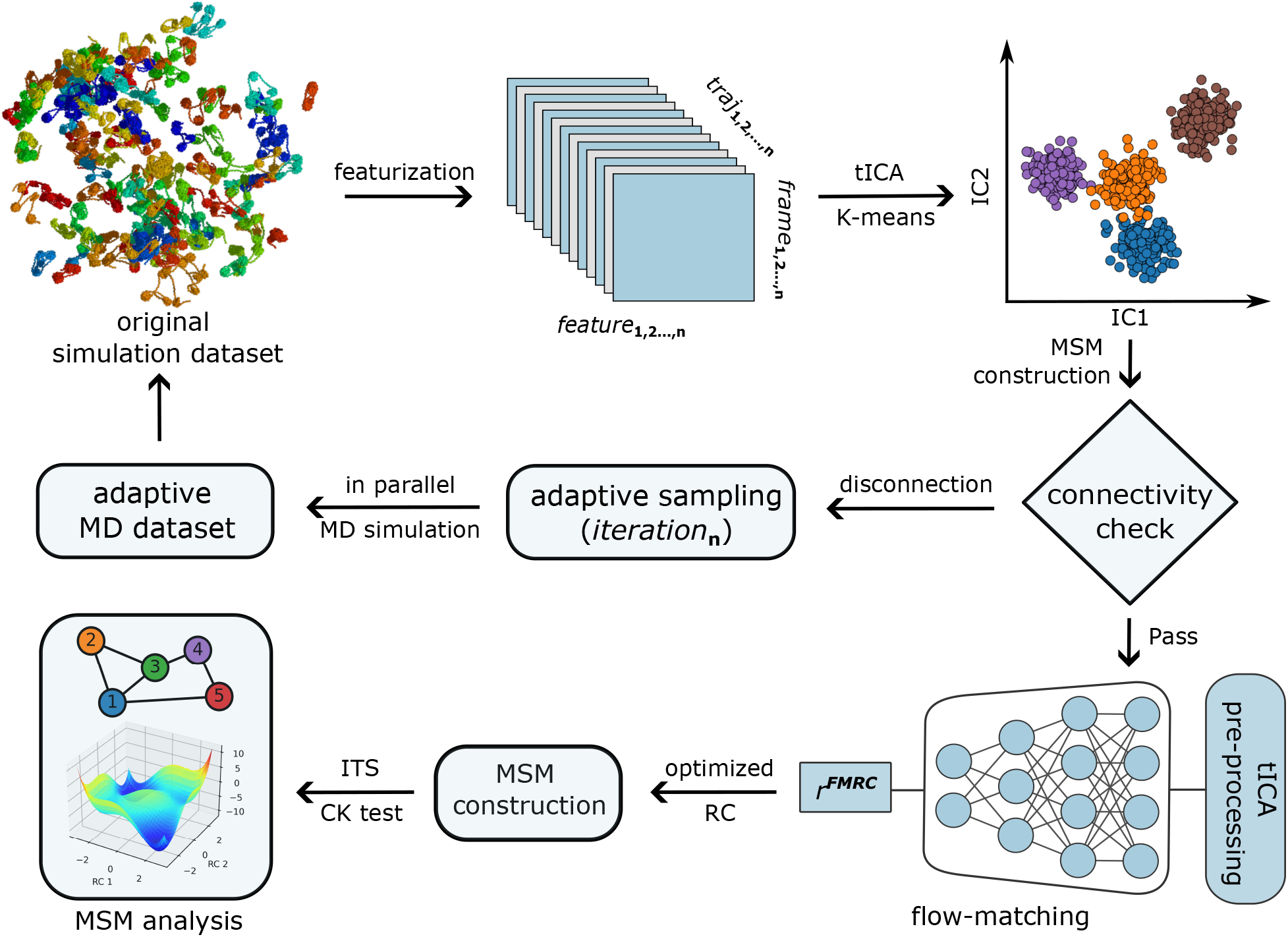
Overall workflow used in this work. Starting from original dataset, we featurize MD trajectories and construct Markov state models. Automated adaptive sampling is performed to resolve potential disconnections in the MSM transition probability matrix built from the original dataset. Downstream reaction-coordinate (RC) optimisation is then applied, and subsequent MSM analysis is carried out once the constructed MSM has been validated using implied timescales (ITS) and Chapman–Kolmogorov (CK) tests.

**FIG. 2.**
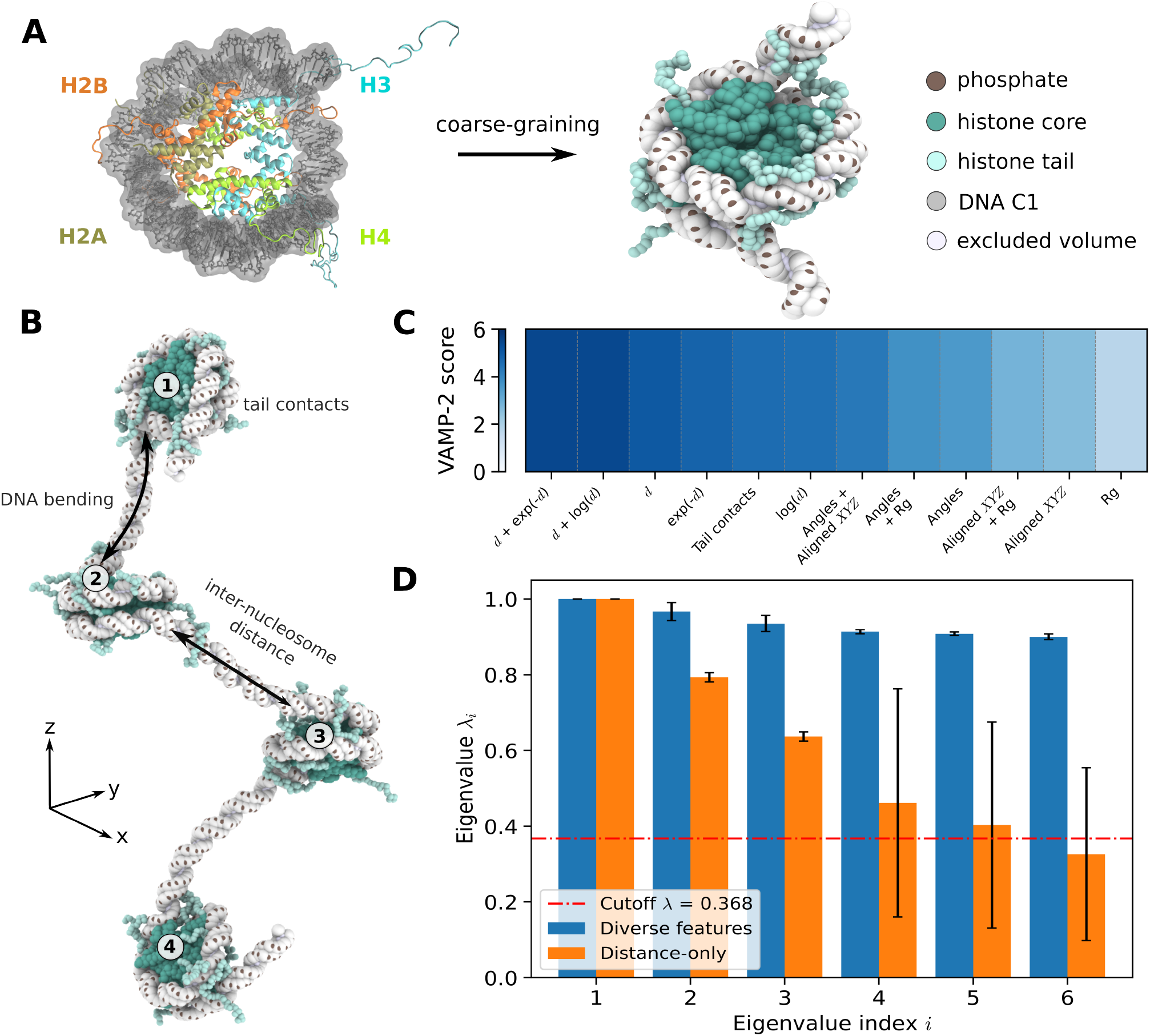
Feature selection for tetra-nucleosome array dynamics using new coarse-grained model. **A**. Mapping of molecular structures to coarse-grained beads in the chemically-specific chromatin model based on the crystal structure of the nucleosome (PDB: 1KX5). **B**. Potential features used to describe tetra-nucleosome folding dynamics. **C**. VAMP-2 scores for each descriptor alone and for combined feature sets. While individual descriptors capture different amounts of dynamics, the combined feature set, including angle, aligned coordinates (RMSD), tail contacts, and raw distances (d) transformed by *e*^−*d*^, log(*d*) consistently achieves the highest VAMP-2 scores. **D**. Comparison of the top six eigenvalues (*λ*_*i*_) of MSMs constructed using distance-only features and diverse feature sets at *τ* = 50 ns. A red dashed line at *λ* = 0.369 indicates the cutoff for eigenmodes with implied timescales shorter than *τ* ^MSM^, used for MSM construction. Eigenvalue uncertainties were estimated by bootstrapping, resampling trajectories with replacement for n = 10 bootstrap replicates.

MD simulations started from distinct configurations selected from D-REMD simulations [45], which yielded a diverse ensemble of chromatin structures under experimentally relevant monovalent salt concentrations of 60–150 mM (see Methods). From these configurations, we launched 320 independent 500 ns unbiased MD simulations, which formed the “original simulation dataset” used in our MSM construction pipeline (Figure 1). Sampling of conformational space was further improved by iterative adaptive sampling (see Methods), in which poorly sampled or disconnected MSM states constructed from existing MD simulations were used as seeds for additional unbiased simulations. In each adaptive iteration, new 500 ns unbiased simulations were launched from these states and merged with the existing data. Several rounds of adaptive sampling were performed for all tetra-nucleosome systems (Table S1), enabling more efficient re-exploration of the conformational space sampled by the existing simulations.

To describe chromatin dynamics in a compact but informative way, we applied Variational Approach for Markov Processes (VAMP)-2 scores [46, 47], a variational metric that quantifies how well features capture slow dynamical processes during chromatin folding. Because our MD simulations used an implicit-solvent coarse-grained model, we initially computed a broad feature set to identify the most informative subset for tetra-nucleosome arrays. Based on previous MSM studies of protein folding [46] and physical intuition about chromatin systems [11, 22, 48], we considered the following descriptor classes (Figure 2B): aligned Cartesian coordinates (e.g., Root Mean Square Deviation or RMSD), distance-based features, tail contacts, flexible torsion angles, and feature combinations (see Methods and SI). Raw distances, *d*, between the centre-of-geometry of two consecutive nucleosomes (i.e., between *i* and *i* + 1; where *i* is the nucleosome index in the array) were also transformed using log(*d*), and *e*^−*d*^ to amplify short-range fluctuations that typically encode slow kinetics and thus highlight important contacts.

Among the individual feature classes, transformed pair-wise inter-nucleosome centre-of-geometry distances yielded the highest VAMP-2 scores, while combined feature sets based on angles and RMSD also showed good performance (Figure 2C). Tail contacts scored highly as well, consistent with the role of histone tails in modulating chromatin folding through electrostatic intermolecular interactions [43, 49]. However, these intrinsically disordered and highly flexible regions are subject to substantial local fluctuations, which are further enhanced by the large integration time step used in coarse-grained simulations. We therefore restricted the final feature set to global chromatin features that directly capture large-scale conformational rearrangements. Collectively, the multi-feature sets used in this work include DNA bending angles, triplet angles between the geometric centres of three consecutive nucleosomes, nucleosome-plane orientations, RMSD, and raw and transformed inter-nucleosome distances (see Methods and SI).

In parallel, we compared MSMs constructed from distance-only features with MSMs constructed from the selected multi-feature representation, using the 30 bp linker system as a benchmark. For the distance-only model, nucleosome indices were symmetrised following Qiu et al. [50]. Compared with the distance-only model, the multi-feature model better captured the slow dynamics of the system. This was reflected by the larger number of slow processes retained above the Markovian cutoff and by faster implied-timescale convergence (Figure 2D and Figure S1). Together, these results underscore the importance of using a sufficiently rich feature space to describe the slow folding dynamics of chromatin fibres.

All trajectories were transformed into VAMP-selected features and projected into a low-dimensional space for clustering and MSM construction. When MSMs of chromatin fibres were constructed directly in tICA space, a small number of leading components captured the dominant slow modes but did not clearly resolve the full complexity of the metastable landscape. Although additional tICA components may contain relevant slow processes, the resulting higher-dimensional representation is difficult to interpret. We therefore employed FMRC [38], an unsupervised deep-learning method that learns compact low-dimensional reaction coordinates for constructing MSMs. Briefly, FMRC learns low-dimensional reaction coordinates by optimising a latent representation that satisfies lumpability and decomposability [51], thereby preserving the time-lagged dynamics encoded in {*x*_*t*_, *x*_*t*+*τ*_} within a reduced latent space. We applied FMRC to tICA-transformed trajectory features, *Ψ*_tICA_, to obtain two-dimensional (2D) latent coordinates, *r* ^FMRC^. MSMs were then constructed and validated using implied-timescale (ITS) analysis and Chapman–Kolmogorov (CK) tests [30, 31]. The resulting microstates were coarse-grained using PCCA+ [52] to define “macrostates”, characterize their conformational ensembles, and quantify kinetic transitions between them. Overall, our pipeline (Figure 1) enabled estimation of chromatin-folding thermodynamics and kinetics from MSMs, providing a basis for quantifying how linker DNA length and sequence shape chromatin structural heterogeneity.

### Linker DNA length controls chromatin polymorphism through distinct free-energy landscape topologies

A central open question in chromatin biophysics is whether chromatin folding can be described by a single, universal energy landscape—analogous to the funnelled landscapes of many globular proteins—or whether its pronounced structural heterogeneity reflects fundamentally different underlying energy landscape topologies. While linker DNA length is known to modulate chromatin structure, compaction, material properties and phase behaviour [9, 43], how these effects are encoded at the level of the a free-energy landscape has remained only partially explored [8, 50]. To address this question, we mapped the folding landscapes of tetra-nucleosome arrays across a broad range of linker DNA lengths using MSMs, allowing us to directly compare landscape topologies as a function of linker geometry. Following the workflow described above, we constructed MSMs for tetra-nucleosome arrays with linker DNA lengths of 15, 20, 22, 25, 30, and 58 bp. After validating each MSM (Figure S2), we computed FESs reweighted by the MSM stationary distributions. Each two-dimensional FES was projected onto the centre-of-geometry distances *d*_13_ and *d*_24_ between nucleosome pairs 1–3 and 2–4, respectively, which capture compaction and stacking in tetra-nucleosome arrays (Figure 3A).

**FIG. 3.**
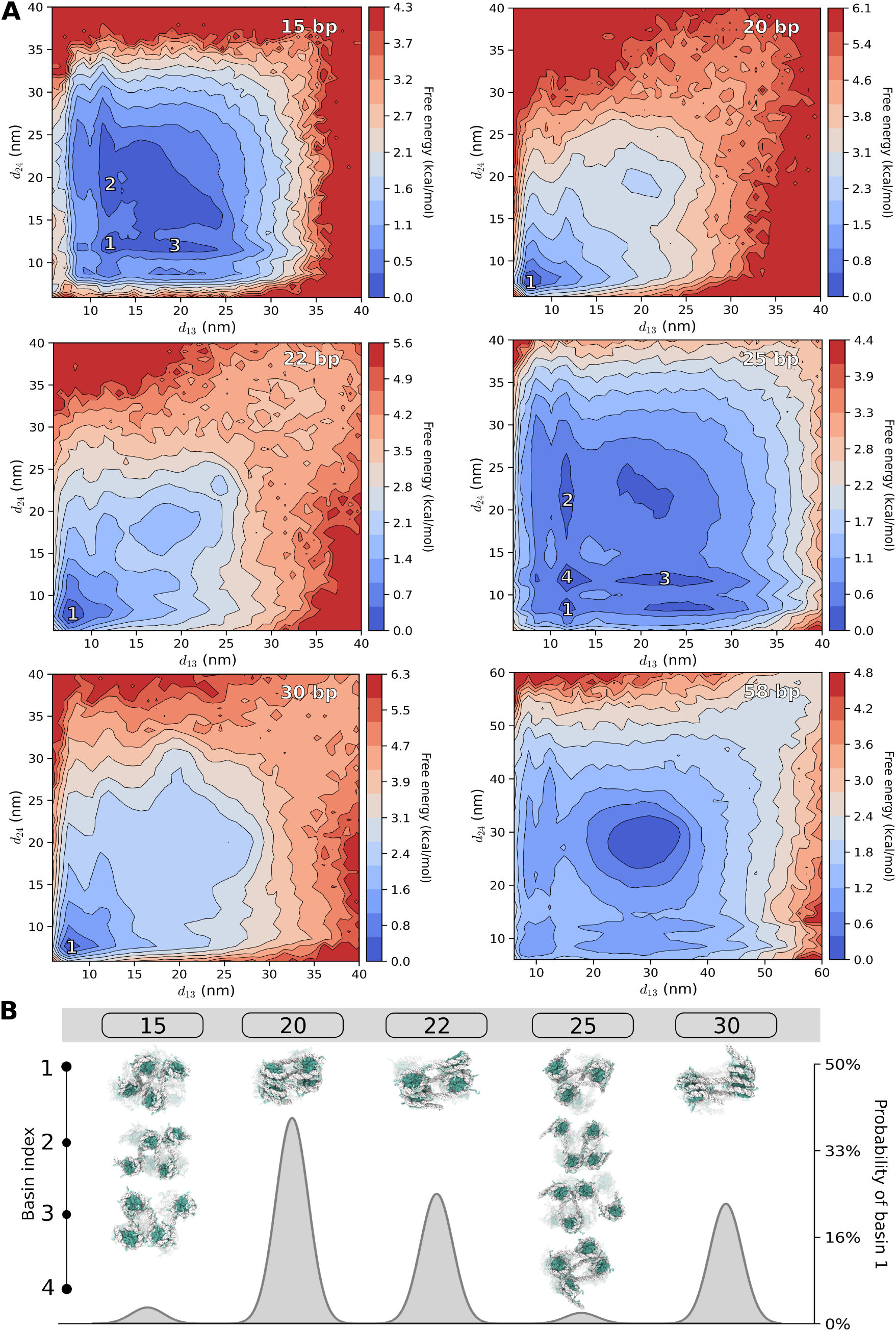
MSM-reweighted free energy landscapes along with inter-nucleosome distance for tetra-nucleosome arrays. **A**. Free-energy landscapes projected onto the centre-of-geometry distances *d*_13_ and *d*_24_ for tetra-nucleosome with linker DNA lengths of 15, 20, 22, 25, 30, and 58 bp. Colour bars are scaled independently for each system, with the upper limit corresponding to the maximum free energy of that system. Landscapes plotted on a common colour scale to facilitate comparison across linker lengths are provided in the Fig S3. **B**. Representative structures from labelled basins and the probability of basin 1 from the FES for near-canonical linker lengths (20, 22, and 30 bp) and non-canonical linker lengths (15 and 25 bp).

The most striking effect of linker DNA length is a transformation of the chromatin free-energy landscape. Depending on linker length, chromatin exhibits funnelled, rugged, or flat landscape topologies, indicating that linker geometry reorganises the number, depth, and connectivity of metastable states and thereby the thermodynamics and kinetics of chromatin folding (Figure 3A). Across all linker lengths, the projected landscapes contain a low-free-energy region corresponding to compact or partially compact conformations and a high-free-energy region at larger inter-nucleosome separations. The total free-energy variation across the (*d*_13_, *d*_24_) projection ranges from ∼ 4.3 to 6.3 kcal/mol, depending on linker length.

These landscape transformations arise from the interplay between two competing physical effects: (i) the geometric frustration introduced by non-integral helical turns of linker DNA, which disrupt the nucleosome orientations required for regular face-to-face contacts, and (ii) the increasing electrostatic and entropic cost of confining longer, highly charged linkers within compact fibre conformations. Face-to-face stacking, in which the flat faces of two nucleosomes are brought into close contact, is the most energetically favourable pairwise nucleosome interaction [9, 11]. In chromatin fibres, this interaction is realised through zig-zag arrangements; i.e., between nucleosomes *i* and *i* + 2. Whether such contacts can form is determined by the rotational phasing imposed by the linker DNA. Because B-form DNA completes one helical turn every ∼10.5 bp [53], linkers close to canonical 10*N* bp values orient successive nucleosomes favourably for this geometry (i.e., parallel nucleosome orientations) [9, 22, 43]. In contrast, 10*N*+5 bp linkers introduce approximately half a helical turn, rotating successive nucleosomes by ∼180^*◦*^ and yielding antiparallel nucleosome orientations, which frustrate face-to-face contact formation [9, 22, 43]. Recovering these interactions then requires linker twisting and bending, generating an elastic deformation penalty that decreases with linker length as the strain is distributed over a longer contour [9]. Linker DNA length therefore sets the balance between the enthalpic gain of nucleosome stacking and the elastic cost required to achieve it, thereby selecting among distinct free-energy landscape topologies.

Linkers close to canonical 10*N* bp values (20, 22, and 30 bp) generate clearly funnelled landscapes characterised by a dominant basin at small *d*_13_ and *d*_24_ values, consistent with parallel nucleosome orientations and compact zig-zag stacking (Figure 3B). For 20 bp, the landscape forms a narrow and steep funnel dominated by basin 1, indicating a strongly preferred stacking geometry. Increasing the linker length to 22 bp broadens the funnel and increases the accessibility of partially open conformations. Mechanistically, the additional 2 bp shift the helical phase by approximately 70^*◦*^, preserving access to stacked configurations while reducing the orientational specificity of face-to-face contacts and lowering the effective barriers separating compact and partially unfolded states.

Extending the linker length from 20 to 30 bp preserves the compact zig-zag corner basin, but the population becomes less concentrated within this single basin and spreads over a broader accessible region of the FES (Figure 3). Thus, compared with 20 bp, the 30 bp linker maintains a compact preferred geometry while allowing greater conformational variability around the stacked state. While longer 10*N* linkers can still support tight face-to-face zig-zag stacking, the longer linker length increases the entropic cost of maintaining a single highly ordered stacked geometry, and the increased DNA charge enhances electrostatic repulsion in compact conformations. As a result, intermediate states become thermodynamically stabilised, producing a wider, shallower funnel without eliminating the dominant compact minimum.

In contrast, intermediate non-canonical 10*N*+5 linkers (15 and 25 bp) generate highly rugged energy landscapes with multiple shallow minima of comparable depth (Figure 3). The antiparallel orientations of successive nucleosome imposed by these 10*N*+5 linkers hinder zig-zag stacking and instead stabilise several competing partially stacked conformations, giving rise to a rugged, multi-basin landscape.

For the long linker of 58 bp (Figure 3A), the landscape becomes comparatively flat and is dominated by a broad, shallow minimum at intermediate inter-nucleosome separations rather than a localised and deep compact corner basin. Although this linker also corresponds to a 10*N*+5 geometry, its greater length allows the imposed rotational mismatch to be absorbed through small torsional and bending deformations distributed along the linker contour. Because these elastic deformation penalties decrease with increasing linker DNA contour length, many distinct conformations have comparable free energy and are separated by low barriers. As a result, structural diversity is magnified (Figure S3) but it arises from continuous fluctuations within a single, weakly structured basin, where many conformations have comparable free energy, rather than from multiple discrete metastable minima.

In summary, chromatin does not possess a single, universal energy landscape. Instead, linker DNA length acts as a control parameter that selects among funnelled, rugged, and flat landscape regimes through geometric frustration of nucleosome stacking. Viewed through this lens, chromatin is best described as a semi-flexible polymer whose degree of frustration is tunable by linker geometry. Its polymorphism therefore reflects not stochastic disorder, but the existence of distinct free-energy landscape regimes with different thermodynamic and kinetic signatures.

### Tetra-nucleosome folding kinetics reveal chromatin is a dynamically reorganising polymer

While the free-energy landscapes described in the previous section characterise the thermodynamic stability of chromatin conformations, they do not by themselves determine how rapidly chromatin folds, rearranges, or interconverts between structures. In complex and frustrated systems, folding kinetics are governed not only by the depth of individual free-energy minima, but also by the connectivity of the landscape, namely the number, accessibility, and barrier heights of pathways linking different conformational states. To quantify how tetra-nucleosome arrays dynamically explore their conformational landscapes, we analysed the MSM transition networks to resolve dominant kinetic pathways, state connectivity, and mean first-passage times (MFPTs) between folded and unfolded macrostates.

We applied PCCA+ [52] to coarse-grain the MSM microstates into five macrostates (Table S2). Rather than behaving like small globular proteins, which typically fold through a small number of dominant pathways into a unique native state and then remain kinetically locked, we find that tetra-nucleosome arrays exhibit multiple inter-macrostate transitions with comparable kinetic weight (Figure 4A). This reflects the nature of chromatin as a semi-flexible polymer with tuneable frustration, whose conformational dynamics are constrained by linker DNA geometry, but are not kinetically frozen. To characterise the structural differences across macrostates, we applied nucleosome– nucleosome contact analysis (see Methods). Specifically, we counted (*i*, *i* + 2) contacts [9, 11]: a larger number of (*i*, *i* + 2) contacts indicates more stacked conformations, whereas fewer contacts correspond to more open or partially folded structures.

**FIG. 4.**
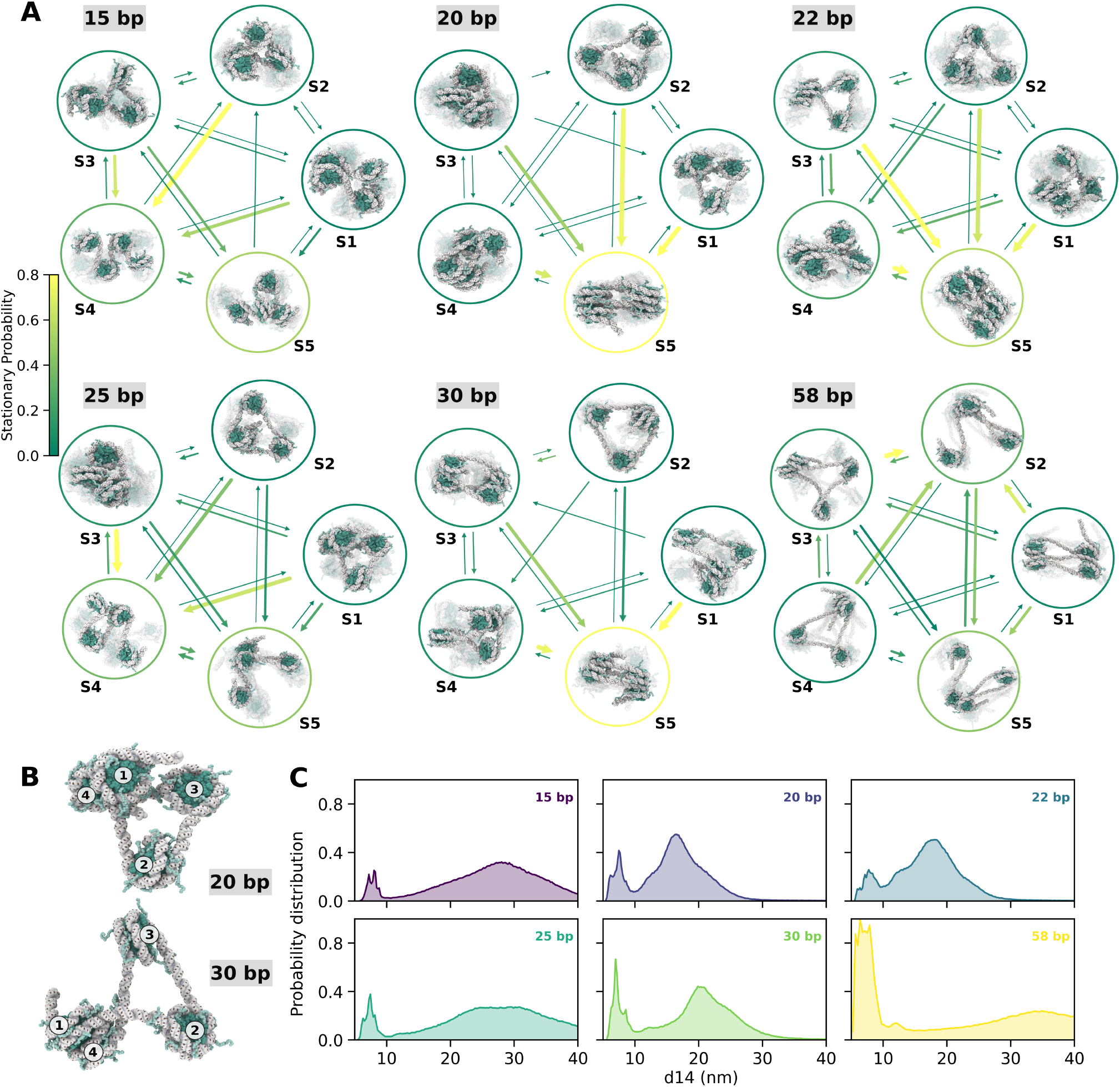
Folding pathways and kinetics between macrostates. **A**. Macrostates obtained via the PCCA+ method. The node edge colour encodes the stationary probability of each macrostate, and the arrow colour encodes the transition probability; both use the same colormap, where lighter colours correspond to higher values. Transitions with very small flux (*T*_*ij*_ ≤ 10^−4^) are filtered out for clarity. The thickness of the arrows is proportional to the transition rate, defined as the inverse mean first-passage time (1*/*MFPT_*ij*_). The thicker arrow refers to a faster transition. **B**. Representative misfolded states at linker lengths of 20 and 30 bp. **C**. Normalised count distributions of conformational ensembles along the centre-of-geometry distance *d*_14_ for linker lengths of 15, 20, 22, 25, 30, and 58 bp.

Consistent with the MSM-reweighted free-energy surfaces, the 20 and 30 bp linkers exhibit a strongly dominant macrostate, S5, which accounts for the majority of the ensemble (Figure 4A and Table S2). Contact analysis shows that S5 is the most compact and internally self-associated state, displaying the largest number of (*i*, *i* + 2) contacts (Figure S5) and therefore the highest degree of intramolecular interaction saturation, or multivalency, within the array. By contrast, the 22 bp system (Figure 4A) exhibits a slightly more heterogeneous stationary distribution: although S5 remains the most populated state, substantial population is also distributed across intermediate macrostates, particularly S4 (Table S2). This redistribution indicates coexistence of multiple metastable conformations, consistent with the broader low-free-energy region observed for 22 bp relative to the more sharply funnelled landscapes of the 20 and 30 bp systems. By contrast, the non-canonical 10*N*+5 arrays, represented by the 15 and 25 bp systems, distribute their stationary probability across multiple macrostates, consistent with a more rugged and frustrated folding landscape. In the 15 bp system, for example, S4 and S5 are populated to similar extents and exhibit similar (*i*, *i* + 2) contact patterns (Table S2 and Figure S5), indicating that multiple conformations coexist with comparable thermodynamic stability. The 58 bp system (Figure 4A) likewise lacks a single overwhelmingly dominant macrostate and instead exhibits several populated metastable states, consistent with a broad, shallow basin that supports a diverse ensemble of irregular conformations.

To compare folding kinetics across linker lengths, we computed mean first-passage times (MFPTs) from less compact macrostates to the most compact macrostate, S5. The estimated MFPTs fall within the microsecond regime of our coarse-grained simulation model and should therefore be interpreted as relative measures of kinetic accessibility at this resolution. For the non-canonical 10*N*+5 linkers, 15 and 25 bp, the rugged landscapes distribute probability across multiple similarly stable basins, allowing frequent exchange among structurally similar folded-like or partially folded conformations. In these systems, transitions toward S5 reflect redistribution within a heterogeneous ensemble rather than relaxation into a uniquely dominant folded state. In contrast, the canonical 10*N* linkers, 20 and 30 bp, are thermodynamically biased toward a compact S5 basin. Their folding kinetics therefore reflect relaxation toward a more strongly selected endpoint, rather than exchange among several comparably populated folded-like states. This distinction shows that linker phasing reshapes not only the equilibrium population of compact states, but also the kinetic organisation of the folding landscape.

For these 20 and 30 bp tetra-nucleosomes, we further identified misfolded states in which nucleosomes 1 and 4 interact first, forming (*i*, *i* + 3) contacts (Figure 4B). We quantified the propensity to form these misfolded states by projecting configurations onto the inter-nucleosome distance *d*_14_. We found a higher probability at short *d*_14_ (≤ 10 nm), which indicates an increased likelihood of entering misfolded (*i*, *i* + 3) contact states (Figure 4C). Notably, this probability is particularly high for the 30 bp linker, providing a mechanistic explanation for its comparatively longer MFPT despite having a highly populated folded basin. Structurally, the (*i*, *i* + 3) contact satisfies locally favourable face-to-face interactions between nucleosomes 1 and 4, but it imposes unfavourable global geometry in terms of linker DNA bending and torsional angle required to reach the (*i*, *i* + 2) stacked zig-zag configuration. Consequently, these states are not thermodynamically competitive, yet they act as kinetic traps that transiently sequester probability flux and delay relaxation into the dominant folded macrostate.

These results show that chromatin behaves as a semi-flexible polymer whose folding kinetics are tuned by linker-length-dependent geometric frustration. For short to intermediate integral 10*N* linkers (20 and 30 bp), most structures occupy a single dominant folded macrostate, yet retain substantial internal dynamics due to accessible kinetic pathways and transient kinetic traps. In contrast, non-canonical 10*N*+5 linkers (15 and 25 bp) are defined by frustrated landscapes with multiple interconverting macrostates of comparable population. Importantly, across all linker lengths, interconversion between folded and partially folded chromatin states is frequent, indicating that compact chromatin fibres remain dynamically reorganising rather than rigid. Together, these findings support a picture in which chromatin is best described as a dynamic polymer, not a static folded structure, capable of continuous internal rearrangement while maintaining structural organisation at the single-fibre level.

### Phase separation reshapes the free energy landscape of chromatin

At physiological salt concentrations, chromatin arrays phase separate into biomolecular condensates whose stability and material properties are governed by dynamic networks of multivalent intermolecular interactions [9, 10, 23]. Notably, arrays with 25 bp linkers form more strongly connected intermolecular networks and more viscoelastic condensates than arrays with 30 bp linkers, reflecting the competition between intermolecular interactions and intra-fibre self-saturation of nucleosome contacts [10, 22]. Furthermore, we previously found that individual 25 bp chromatin fibres undergo conformational expansion upon phase separation [22]. This observation suggests that the free-energy landscape of a chromatin fibre may depend on its condensed environment and the intermolecular interactions it experiences therein.

To test this idea, we performed direct-coexistence simulations of 25 bp tetra-nucleosome chromatin condensates using OpenCGChromatin, as done previously [22]. Briefly, 81 tetra-nucleosome arrays were placed in an elongated periodic simulation box, allowing a chromatin-rich condensate to coexist with dilute phases separated by planar interfaces. The simulations were performed at room temperature and *λ*_*D*_ = 1.0 nm, corresponding to approximately 100 mM monovalent salt. This direct-coexistence setup allows chromatin molecules to exchange freely between the dense and dilute phases, enabling us to analyse the conformational ensemble of individual tetra-nucleosome arrays inside the condensate.

Using 10 *µ*s trajectories, we constructed and validated MSMs in the *r* ^FMRC^ space (Figure S6), computed the corresponding free-energy landscapes and projected each tetra-nucleosome onto the same inter-nucleosome distances used above, *d*_13_ and *d*_24_. We then compared the landscape obtained from tetra-nucleosomes in the bulk of the condensed phase (54 tetra-nucleosomes) with that obtained from all tetra-nucleosomes in the simulation box, spanning the bulk, interface, and dilute phases.

Remarkably, the free-energy landscape sampled by chromatin arrays within the condensate differs substantially from that of isolated fibres (Figures 5 versus 3A). Whereas the isolated 25 bp tetra-nucleosome system exhibits a rugged landscape with several well-defined minima corresponding to distinct compact and partially compact conformations, the condensed-phase landscape is markedly smoother and dominated by states at larger inter-nucleosome separations. Upon phase separation, the compact minima that characterise the isolated-fibre landscape are substantially depleted, and statistical weight is redistributed toward a broad ensemble of partially expanded conformations. Consistently, restricting the analysis to chromatin arrays buried within the condensate interior (i.e., excluding fibres at the interface and in the dilute phase) further accentuates this shift (Figure 5 left versus right), indicating that the local condensate environment selectively stabilises expanded chromatin states.

**FIG. 5.**
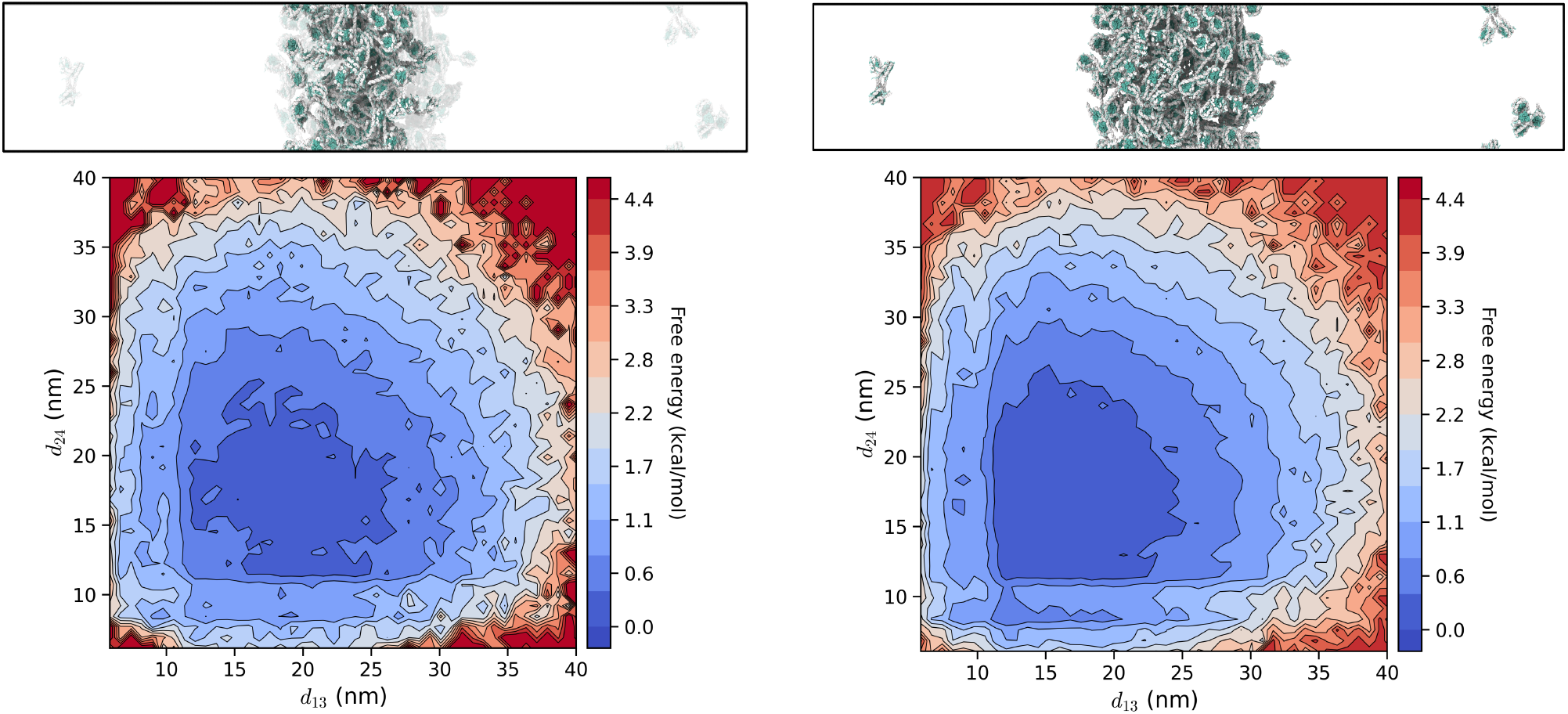
Free-energy landscapes of the 25 bp tetra-nucleosome system in a chromatin condensate (*N* = 81 tetra-nucleosome arrays total). The left panel shows tetra-nucleosome arrays in the condensed phase (*N* = 54), and the right panel shows the full ensemble (*N* = 81). Chromatin fibres in the condensed phase were selected based on the density profile analysis described in Methods.

These changes to the free-energy landscape of chromatin are physically consistent with the intermolecular network organisation of chromatin condensates [22]. When chromatin adopts compact conformations its nucleosome faces and histone tails are internally self-saturated through intra-fibre contacts, which limits their accessibility for intermolecular interactions. Partial fibre expansion within the condensate is important because it exposes nucleosome surfaces and histone tails, increasing the effective valency of chromatin and allowing it participate in the formation of the inter-molecular interaction network that stabilises the dense phase.

These results demonstrate that phase separation reshapes the free-energy landscape of chromatin by altering the statistical weights of the different accessible microstates, favouring the partially expanded, disordered states that maximise the density of inter-molecular connections and the enthalpy gain upon phase separation.

### Single base-pair variations in linker DNA generate a continuum of chromatin free-energy landscape and phase separation regimes

Linker DNA length controls both inter-nucleosome spacing and orientation. Even a 1 bp change can strongly perturb chromatin folding [9]. In the chromatin fibre, this phase shift alters how the linker DNA can bend and twist to accommodate a given packing geometry and whether nucleosome–nucleosome faces can align to support stable zig-zag stacking. To quantify the impact of single-bp perturbations, we also computed free-energy landscapes projected onto the *d*_13_ and *d*_24_ for linker lengths of 26, 27, 28 and 29 bp after MSM validation (Figures 6A and S7).

**FIG. 6.**
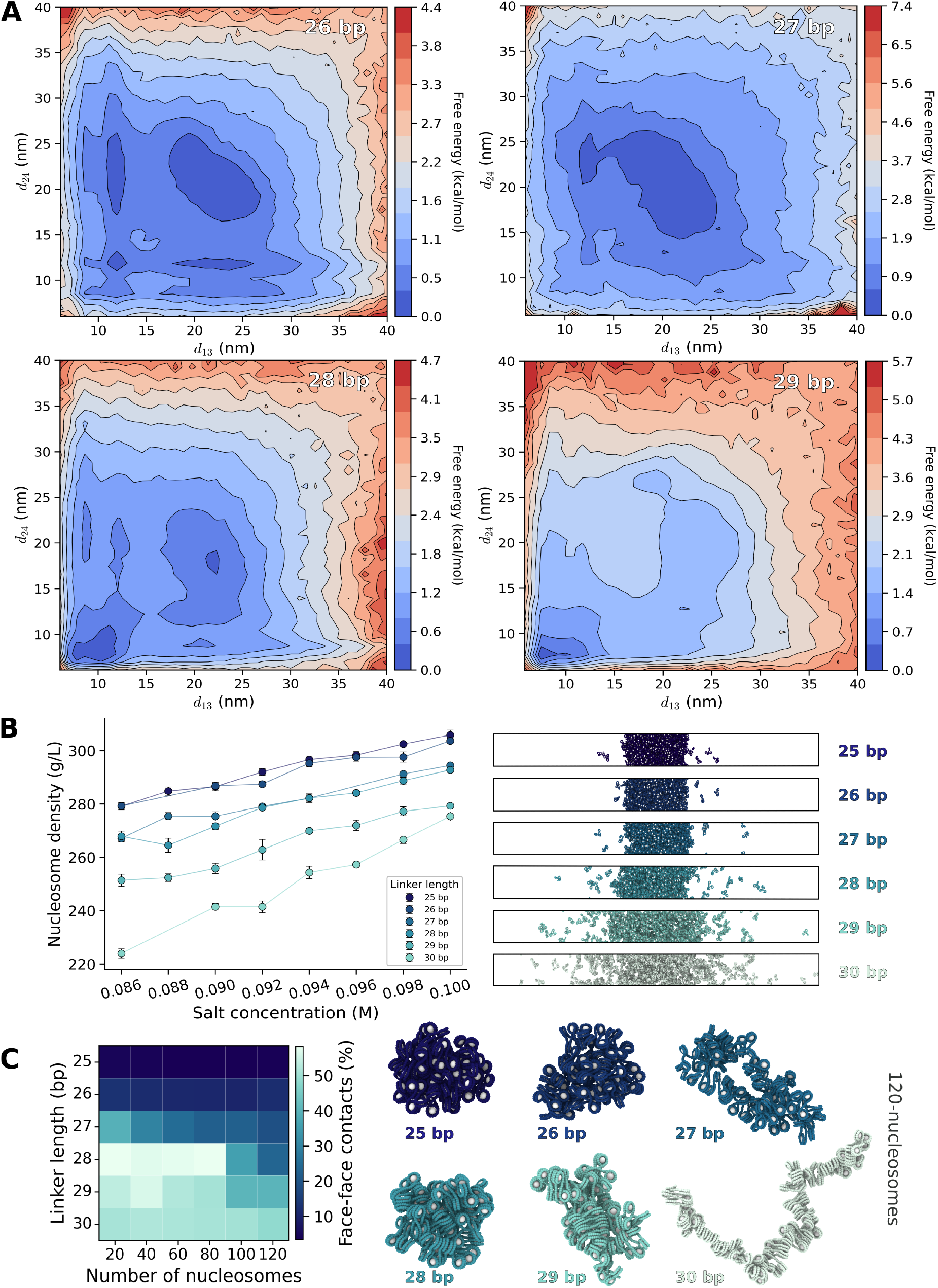
Effects of single-base-pair linker increments on tetra-nucleosome free-energy landscapes, phase separation, and chromatin array structure. **A**. Free-energy landscapes projected onto centre-of-geometry distances *d*_13_ and *d*_24_ for tetra-nucleosome arrays with 26– 29 bp linker DNA. Landscapes for 25 and 30 bp are shown in Fig. 3A. Colour bars are scaled independently; common-scale landscapes are provided in Fig. S3. **B**. Condensed-phase density profiles of tetra-nucleosome systems with 25–30 bp linker DNA as a function of salt concentration. Representative condensate snapshots at 0.086 M salt are shown, with histone cores in white and linker/nucleosomal DNA in the same colour. **C**. Average percentage of face-to-face contacts for *N*-nucleosome arrays with 25–30 bp linker DNA. Representative 120-nucleosome structures are shown on the right and rendered as in Fig. 6B.

The resulting landscapes reveal that chromatin does not transition between a small number of discrete landscape classes. Strikingly, linker length non-linearly reshapes the landscape, producing a spectrum of intermediate landscape regimes between rugged and funnelled limits (Figures 3A and 6A). At 25 bp, the free-energy surface is multi-basin, indicating an ensemble in which several conformational states are thermodynamically competitive. Increasing the linker length to 26 and 27 bp progressively smooths the landscape, with initially distinct basins merging into a broader low-free-energy region. This behaviour suggests an intermediate regime in which both zig-zag–like conformations and disordered structures remain accessible. A qualitative transition occurs near 28 bp, where compact conformations with smaller *d*_13_ and *d*_24_ first become thermodynamically favourable. As the linker length further approaches the canonical 10*N* value, the landscape increasingly narrows toward compact states, and by 30 bp it is dominated by a well-defined basin consistent with stable face-to-face nucleosome stacking. Thus, single-base-pair linker changes tune the balance between frustrated multi-basin folding and a funnelled compact zig-zag state.

These single-fibre landscapes are consistent with the linker-length-dependent phase behaviour observed in direct coexistence simulations. Using our minimal chromatin model [11] (see Methods and Figure S8), we performed additional direct coexistence simulations of condensates assembled by tetra-nucleosome arrays with linker lengths spanning the 25–30 bp series (see SI). In Figure 6B, we show the equilibrium densities of the condensates for the different linker lengths as a function of salt concentration. Consistent with experiments, regardless of linker length, condensate density increases with salt, as expected from the dominant effect of DNA–DNA repulsion in opposing chromatin compaction and phase separation [9, 10]. More importantly, as observed experimentally for 12-nucleosome arrays [9], the equilibrium densities of the condensates decrease non-linearly as linker length increases from 25 to 30 bp (Figure 6B). This trend can be rationalised by the redistribution of nucleosome face-to-face interactions between intra- and intermolecular contacts. Linkers near 10*N*+5 favour heterogeneous, less internally saturated fibre conformations, leaving nucleosome interaction surfaces available for intermolecular association and thereby promoting phase separation. In contrast, linkers approaching 10*N* increasingly stabilise intramolecular zig-zag stacking, which satisfies these same interactions within individual fibres and reduces the driving force for condensation.

While these free-energy and phase-behaviour analyses reveal how linker length modulates the equilibrium balance between disordered and compact fibre states, they do not directly identify the structural coordinates governing the slow folding dynamics. To address this, we examined the two slowest MSM modes, *ψ*_2_ and *ψ*_3_, for each linker length and computed the maximum absolute correlation between these modes and our diverse feature sets. Notably, the linker torsion angle, DNA RMSD, and nucleosome-core RMSD (Figure S9) are the top three features with high correlation values for all linker lengths. More interestingly, these three features exhibit a specific pattern across this 1 bp series. In particular, the torsion angle contribution decreases from 25 to 27 bp and remains nearly unchanged at 28 bp, but increases markedly at 29 bp and peaks at 30 bp. DNA RMSD shows a similar pattern: correlations remain comparable from 25 to 28 bp and then rise sharply at 29–30 bp. In contrast, nucleosome–core RMSD exhibits the opposite trend, with the largest correlation at 25 bp and a consistent decrease as the linker length increases.

This suggests a competition between two limiting mechanisms for the slow relaxation. For short, non-canonical linkers (25–26 bp), the rotations and higher torsional stiffness of DNA frustrate the simultaneous formation of symmetric, compact stacking contacts. Consequently, the slow modes are dominated by rearrangements of nucleosome cores across multiple shallow basins. The relatively modest barriers between these basins imply rapid interconversion, so adding only 1–3 bp is sufficient to merge unfolded and partially folded ensembles into a more coherent zig-zag-like basin. Near the canonical cases (29–30 bp), the linker is long enough and the rotational setting favourable enough to stabilise compact stacking, so the bottleneck for interconversion shifts toward DNA-mediated degrees of freedom, where the fibre must twist and deform the linkers to disrupt and re-establish these contacts. Accordingly, torsion and DNA conformational changes become the dominant contributors to the slow MSM modes, while nucleosome–core rearrangements play a progressively smaller role.

To test whether the linker-length-dependent reshaping of the free-energy landscape persists beyond tetra-nucleosomes, we turned to our minimal chromatin coarse-grained model [11] to simulate chromatin arrays containing up to 120 nucleosomes. Because extending the adaptive sampling of MSM framework used for tetra-nucleosome arrays to fibres of this size would require prohibitively extensive sampling, we did not attempt to reconstruct the corresponding free-energy landscapes. Instead, we performed temperature molecular dynamics (T-REMD) simulations (see Methods) of the minimal chromatin arrays containing *N*_*c*_ = (20, 40, 60, 80, 100, 120) nucleosomes, where *N*_*c*_ denotes the number of nucleosomes in the fibre, for linker lengths of 25, 26, 27, 28, 29 and 30 bp. From these simulations, we quantified the fraction of face-to-face nucleosome contacts as a structural proxy for the degree of landscape funnelling inferred from the tetra-nucleosome analysis (Figure 6C). Consistent with the free-energy landscapes obtained for tetra-nucleosome arrays at near-atomistic resolution (Figures 3 and 6A), longer fibres exhibit a pronounced dependence of nucleosome stacking on linker length. Arrays with 25–27 bp linkers form relatively few face-to-face contacts across all fibre lengths, whereas linker lengths of 28–30 bp support substantially greater zig-zag contact formation. Notably, the transition observed in the tetra-nucleosome landscapes near 28 bp reappears in the longer fibres as a sharp increase in stacking contacts. The emergence of this transition in fibres spanning up to 120 nucleosomes suggests that the local geometric constraints imposed by linker DNA propagate cooperatively through the fibre, amplifying the structural consequences of single-base-pair changes in nucleosome spacing. Moreover, this effect becomes increasingly pronounced with fibre length, indicating that small linker-length-dependent differences in local folding are progressively amplified in larger chromatin arrays. Representative structures of 120-nucleosome fibres further illustrate this behaviour (Figure 6C), revealing marked differences in fibre organisation and stacking patterns arising from single-base-pair variations in linker length.

Together, these results reveal that single-base-pair changes in linker length reshape chromatin behaviour across multiple scales. At the level of individual tetra-nucleosome arrays, they alter the topology of the free-energy landscape and the dominant slow modes of structural relaxation. These changes are accompanied by differences in condensate density and phase-separation propensity, and their structural signatures persist in extended chromatin fibres, where they become increasingly amplified with array length. Importantly, our results show that rather than populating a small number of discrete structural states, chromatin spans a continuum of linker-length-dependent landscape regimes whose consequences propagate from local linker-DNA geometry to fibre architecture and collective condensate behaviour.

### DNA sequence-dependent remodelling of chromatin free-energy landscapes

In addition to linker length, the sequence of the linker DNA can remodel chromatin folding by tuning the local mechanical response of the linker, including bending, twist elasticity, and the amplitude of thermal fluctuations. We therefore compared three linker sequences, a mixed-composition linker (`ACTG`), `polyA` and `polyAT`, at two representative linker lengths, for a 10*N*+5 linker (25 bp) and a 10*N* linker (30 bp). Among these DNA sequences, `polyA` is the stiffest as it exhibits the highest DNA deformation energy, while `polyAT` is the softest [54].

For 25 bp linkers, changing the DNA sequence to either `polyA` or `polyAT` substantially alters both the shape and the depth of the minima in the free-energy landscapes (Figure 7A). As discussed above, the 25 bp linker length corresponds to approximately 1.5 turns of DNA (10*N*+5), which results in face-to-face nucleosome stacking not being possible without mechanically deforming the linker DNA. As a result, the energetic cost of deforming the linker DNA plays a major role in determining the structures the chromatin fibre can adopt. Sequences that are more deformable can more easily accommodate these distortions, allowing nucleosomes to approach and form partial stacking contacts, whereas stiffer sequences penalise these adjustments and favour more disordered conformations.

**FIG. 7.**
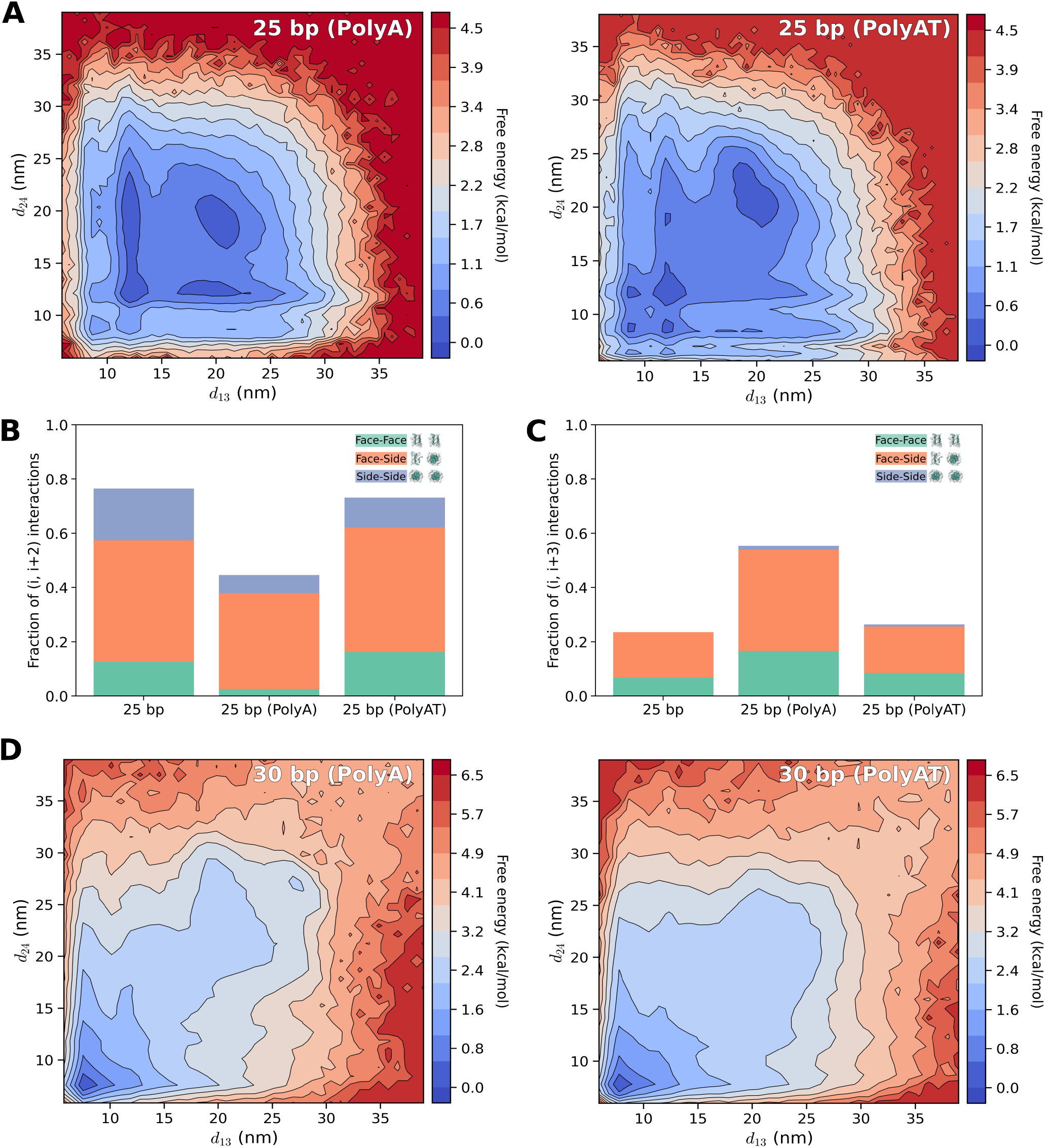
DNA sequence-dependent free-energy landscapes. **A**. Free-energy landscapes along the centre-of-geometry distances *d*_13_ and *d*_24_ for a 25 bp linker and two DNA sequences: `polyA` and `polyAT`. **B**. (*i*, *i* + 2) inter-nucleosome contact analysis for conformational ensembles extracted from local basins, with basin-resolved contact fractions averaged using basin probabilities from the FES. **C**. (*i*, *i* + 3) inter-nucleosome contact analysis for conformational ensembles extracted from local basins, with basin-resolved contact fractions averaged using basin probabilities from the FES. **D**. Free-energy landscapes along the centre-of-geometry distances *d*_13_ and *d*_24_ for a 30 bp linker and DNA sequences: `polyA`, `polyAT`.

To connect these energy landscape differences to specific chromatin structural changes, we quantified inter-nucleosome contact patterns within the dominant basins for the different 25 bp systems (Figure 7B). For the mixed `ACTG` linker, contacts are dominated by the (*i*, *i* + 2) pair, which corresponds to the zig-zag stacking register, although, in agreement with the structural disorder of this systems discussed above, most interactions are face-to-side rather than face-to-face. A smaller fraction of face-to-face stacking contacts is nevertheless present. For the stiffer `polyA` linker, the fraction of (*i*, *i* + 2) face-to-face contacts is strongly reduced, indicating that the linker resists the bending and torsional deformation required to bring nucleosome faces into the stacking register. Instead, the fibre samples more alternative interaction patterns, including an increased population of (*i*, *i* +3) contacts. Consistently, the more deformable `polyAT` linker promotes (*i*, *i* + 2) stacking contacts, including face-to-face interactions.

In contrast, for 30 bp linkers the free-energy landscapes are nearly indistinguishable across `ACTG`, `polyA`, and `polyAT` (Figure 7D), each displaying a single dominant minimum corresponding to compact zig-zag conformations. At this canonical linker length, the linker DNA completes an integer number of helical turns (10*N*), aligning nucleosome faces in the preferred stacking register. Because the geometry already favours face-to-face nucleosome interactions, compact stacking can form without requiring significant deformation of the linker DNA. In this regime, the energetic gain from nucleosome stacking dominates over modest sequence-dependent differences in DNA mechanics, rendering the overall energy landscape largely sequence-independent.

This contrast highlights that sequence-dependent DNA mechanics become most important when the chromatin fibre has linker lengths close to 10*N*+5 that frustrate face-to-face stacking. Under these conditions, small differences in linker deformability reshape the accessibility of stacked conformations and reorganise the free-energy landscape. By contrast, when the linker length naturally supports compact face-to-face stacking, as for canonical 10*N* linkers, sequence effects mainly modulate fluctuations within the dominant stacked state rather than altering the overall landscape.

## DISCUSSION

By mapping the folding landscapes of tetra-nucleosome arrays with adaptive MSMs built on near-atomistic coarse-grained simulations, we find that chromatin is characterized by a continuum of free-energy landscape topologies, ranging from funnelled, to rugged, to flat isosurfaces. This finding fundamentally reframes chromatin polymorphism: the structural heterogeneity of chromatin fibres does not reflect thermal fluctuations within a single dominant basin, but the coexistence of multiple metastable states and, across linker geometries, movement along a continuous spectrum of landscape topologies. Therefore, no single structure (e.g., zig-zag versus solenoid) characterises a chromatin fibre. The more fundamental description is the free-energy landscape, from which the observable ensemble follows the accessible states from its basins, their relative stability from basin depths, and their interconversion kinetics from the barriers between them.

The parameters that determine different free-energy landscape topologies are the linker DNA length and sequence, through the geometric frustration they impose on the fibre conformation. For near-canonical 10*N* linkers, the fibre internally self-saturates its strongest face-to-face nucleosome contacts, producing a funnelled landscape with a dominant compact zig-zag basin. Because face-to-face stacking forms without appreciable linker deformation, these landscapes are largely sequence-independent. At 10*N*+5 linkers, the extra half-helical turn of linker DNA rotates successive nucleosomes by roughly 180^*◦*^ relative to one another, frustrating symmetric face-to-face stacking and forcing the fibre to choose among discrete, only partially satisfied configurations, thereby generating a rugged landscape of competing minima. Here, sequence becomes important because deformation of the linker DNA is required to recover favourable stacking interactions: a soft sequence such as `polyAT` facilitates this deformation and enables (*i*, *i* + 2) face-to-face contacts, whereas a stiff sequence such as `polyA` resists deformation and shifts the ensemble toward disordered, alternative-contact configurations [54]. For long linkers, frustration is relieved by distributing torsional and bending deformation along the extended contour, flattening the landscape into a broad, entropically rich basin and diminishing the influence of sequence.

This control operates with single-base-pair precision. Increasing the linker length from 25 to 30 bp one base pair at a time continuously reshapes the free-energy landscape, generating a spectrum of topologies that spans from rugged, multi-basin landscapes (25–26 bp) through an intermediate broad-basin regime (27 bp) to funnelled landscapes dominated by compact zig-zag conformations (28–30 bp). The accompanying crossover in the slowest kinetic modes—from nucleosome-core rearrangements among competing shallow minima (25–26 bp) to linker-DNA torsion and bending (28– 30 bp)—demonstrates that this transition reflects a fundamental reorganisation of the dominant relaxation pathways. These landscape transformations provide a mechanistic explanation for the non-linear modulation of chromatin phase behaviour by single-base-pair changes in linker length observed experimentally [9]. Rugged landscapes (25–26 bp) disfavour intramolecular self-saturation, leaving nucleosome faces and histone tails available for intermolecular association and thereby promoting the connectivity of the network that drives phase separation. In contrast, funnelled landscapes (28–30 bp) increasingly satisfy these interactions within individual fibres, reducing intermolecular connectivity and suppressing phase separation. Therefore, the non-linear phase behaviour observed across this 1-bp linker series experimentally [9], emerges directly from the spectrum of free-energy landscape topologies of chromatin.

A central conceptual implication of our work concerns the nature of the energy landscape itself. Free-energy landscapes are generally treated as intrinsic properties of molecules, defined in dilute solution and independent of the collective state of the system. Our results challenge this view. Our direct-coexistence simulations show that the landscape of a tetra-nucleosome inside a condensate differs from that of the isolated fibre, with statistical weight shifted from the most compact corner basins toward expanded, disordered conformations. The free-energy landscape is thus not a fixed, intrinsic property of the molecule but an emergent, environment-dependent one. The condensate redistributes population toward partially expanded states because these maximise the density of intermolecular contacts and the enthalpic gain of the network. The coupling between landscape and phase separation therefore runs in both directions. That is, disordered single-fibre states promote phase separation, and phase separation in turn stabilises them. Thus, the conformational properties of a chromatin fibre are co-determined by the phase in which the fibre resides.

Although our free-energy landscapes were reconstructed for tetra-nucleosome arrays at near-atomistic resolution, the corresponding linker-length-dependent stacking trends persist in minimal chromatin fibres containing up to 120 nucleosomes. Nevertheless, longer arrays at near-atomistic resolution may introduce additional long-range frustration and landscape features beyond those mapped here. The systems studied also do not include post-translational modifications, binding of additional proteins, nucleosome breathing, intra-array linker DNA variability, or explicit solvent and ions, each of which is expected to reshape the free-energy landscapes of chromatin. We anticipate, however, that incorporating these additional sources of heterogeneity would enrich rather than overturn the picture developed here, expanding the repertoire of accessible landscape topologies and reinforcing the view of chromatin as a polymer whose polymorphism is encoded in a tunable free-energy landscape.

Taken together, our results motivate a conceptual revision of how chromatin fibre structure and its link to phase separation should be understood. The folding-funnel paradigm, so successful for many globular proteins, does not transfer to chromatin. Chromatin is better viewed as a frustrated heteropolymer spanning a continuum of landscape topologies. This continuum reconciles a long-standing tension in the field—the ordered zig-zag fibres observed in reconstituted arrays in vitro [41, 42] and the disordered chromatin organisation observed in cells [3–5, 7] are not competing models, but different regimes of a single landscape continuum, selected by parameters like linker geometry, chemical heterogeneity, and solution conditions. It also decouples compaction from accessibility: building on the finding that compact, irregularly structured chromatin remains searchable by DNA-binding factors [11]. Our kinetics show that even ordered, funnelled 10*N* fibres interconvert on the microsecond timescale, so chromatin compaction does not unequivocally imply static DNA inaccessibility.

Linker DNA length varies across species, cell types, and genomic regions and is actively regulated by chromatin-binding proteins [55, 56]. Therefore, we suggest that such variation reshapes chromatin not by selecting individual structures, but by tuning the energy landscape from which those structures emerge: a change in linker geometry alters the topology of the landscape itself and, with it, the ensemble of accessible conformations, their interconversion dynamics, and their propensity to phase separate. More broadly, we show that chromatin organisation is governed by free-energy landscapes that couple across scales, linking base-pair DNA mechanics to mesoscale compartmentalization—capturing not only which conformations chromatin populates, but how they interconvert and self-assemble into mesoscale compartments such as condensates.

## MATERIALS AND METHODS

### Coarse-grained models and simulation setup

#### Chemically-specific coarse-grained model

The structured histone core of each octamer is represented by an elastic network model (ENM), in which each bead corresponds to one amino acid and preserves its atomistic hydrophobicity and charge. Intrinsically disordered histone tails are modelled as flexible polymers connected only by harmonic bonds. DNA mechanics are described by sequence-dependent bond and angle potentials derived from the CGeNArate framework [57]. Detailed model parameters can be found in Russell et al. [22]

Simulations of single-fibre systems were performed at Debye length of 0.8 nm and a temperature of 298 K. Each nucleosome contains 147 base pairs (bp) of DNA, and the inner 127 bp of nucleosomal DNA were permanently attached to the histone core in all simulations. The linker length was set to be 15, 20, 22, 25, 26, 27, 28, 29, 30 and 58 bp with linker DNA sequence of ‘`ACTG`’. Regarding to cases of 25 and 30 bp, we also replaced the linker DNA sequence to ‘`AAAA`’ and ‘`ATAT`’ to explore the effect of DNA rigidity on the free energy surface. The simulations were all performed in OpenMM [44] (version 8.1.2). `LangevinMiddleIntegrator` was employed with timestep 10 fs and friction coefficient *γ* = 0.01 ps^−1^. The parallel simulations script implemented CUDA Multi-Process Service (MPS) so as to achieve better simulation performance. The tetra-nucleosome condensate simulation with a 25 bp linker used trajectory from a previous study [22]. 2000 frames were saved from the last 10 *µ*s of 20 *µ*s trajectory and used in this work.

#### Minimal coarse-grained model

The minimal model provides a highly coarse-grained representation while retaining the key features needed to reproduce chromatin structural ensembles obtained from the chemically specific model (Figure S8). In this representation, each histone octamer is reduced to a single bead, which preserves the overall nucleosome geometry, the arrangement of DNA around the histone core, and the effective nucleosome–nucleosome interactions parameterised from the chemically specific model. This level of coarse-graining enables the extensive sampling required to construct full phase diagrams, which would be computationally prohibitive using higher-resolution simulations. Further details of the minimal model are provided in Farr et al., [11] and the chromatin condensate simulations using minimal CG model is described in the Supplementary Information.

### Debye length Hamiltonian replica exchange

We used Debye length replica exchange molecular dynamics (D-REMD), a Hamiltonian REMD scheme in which the Debye length was varied across replicas to promote conformational sampling of the chromatin model. All replicas are maintained at the same temperature, but each is assigned a different value of *λ*_*D*_, thereby modulating the strength of electrostatic interactions. Exchanges between two neighbouring replicas *i* and *i* + 1 were proposed every 10 ps and evaluated using the Metropolis rule:

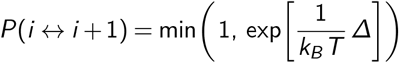

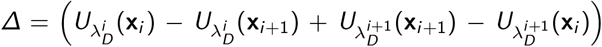

where **x**_*i*_ represents the coordinates of the *i* ^th^ replica, and 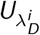 is the potential energy at Debye length 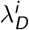. Debye lengths were distributed according to an exponential ladder, 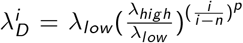, where *n* is the total number of replicas and *p* = 1.25 controls the spacing. The ladder was tuned to achieve an exchange acceptance rate of approximately 20–40 % between neighbouring replicas. We applied D-REMD to a tetra-nucleosome array with linker lengths of 15, 20, 22, 25, 26, 27, 28, 29, 30 and 58 bp as well as the case of using `polyA` and `polyAT` as linker DNA sequence. We generated 16 replicas spanning Debye length from 0.8 (low salt: 60 mM) to 1.3 nm (high salt: 150 mM) at 298 K. A total of 120000 exchange attempts were performed at 10 ps intervals, yielding 1.2 *µ*s of sampling for each replica. The simulations were run in OpenMM using the openmmtools `ReplicaExchangeSampler` with `LangevinDynamicsMove` [44, 58].

### Temperature replica exchange molecular dynamics

We used temperature replica exchange molecular dynamics (T-REMD) of chromatin arrays represented with our minimal coarse-grained model [11] with replicas spanning a temperature range of 300–600 K. Depending on the system size we used an appropriate number of replicas to give exchange probabilities close to 0.3. The simulations were run for at least 200 million timesteps, with T-REMD exchanges attempted every 100 timesteps. Coordinate snapshots were recorded every 100 thousand timesteps. The first half of the trajectories were neglected from the analysis. Only the 300 K replicas were used for analysis.

### Adaptive sampling

The adaptive sampling scheme was built upon Zhang et al [25]. Specifically, in each adaptive sampling round, the tICA lag time was set to 100 ns, and the leading eigenvectors contributing 95% of the kinetic variance were retained [59]. The configurations were then clustered into discrete states, or microstates, using K-means. The number of microstates used in each round was chosen using a heuristic approach (https://github.com/Acellera/htmd/blob/master/htmd/adaptive/adaptivebandit.py). The MSM lag time was set to 250 ns, and microstates with lower counts were assigned a higher probability of being selected as seeds for adaptive sampling. The resulting microstates were then coarse-grained into 20 macrostates using PCCA+ [52]. In each adaptive sampling round, 20 seed positions were selected, and each seed configuration was propagated for 500 ns using the same simulation setup described above. Table S1 summarizes the number of adaptive sampling rounds performed for each tetra-nucleosome system and the total number of trajectories used for MSM construction.

### Feature selection and evaluation

Features were written to `COLVAR` files using PLUMED version 2.9.2 [60, 61]. Input files were generated by an in-house Python script. In total, 2720 features were generated for each tetra-nucleosome array and grouped into raw distances, transformed distances, RMSD, and angles. Specifically, the nucleosome globular domain (724 beads) was partitioned into 15 segments. Inter-segment distances were computed to obtain inter-nucleosome distances (*d*_12_, *d*_13_, *d*_14_, *d*_23_, *d*_24_, *d*_34_), thereby enriching the feature set. An exponential transformation, *e*^−*d*^, was applied to the raw distances to emphasize short-range fluctuations.

We further incorporated features describing DNA structure and nucleosome arrangements, including linker-DNA bending angles, torsion angles (*ϕ*) defined by the planes of nucleosomes 1–2–3 and 2–3–4, and the corresponding angles between these planes (with nucleosomes 2 and 3 treated as vertices, respectively). We also computed RMSD values for the full DNA chain and for the four nucleosome cores relative to representative structures obtained from clustering. Local interactions were quantified via contacts between histone tails and nearby DNA segments or globular domains. These feature groups were evaluated using the `VAMP-2` score function in Deeptime [59, 62]. Detailed feature descriptions, mathematical definitions, and evaluations are provided in the Supplementary Information.

### Reaction coordinate learning and Markov state modelling

Regarding reaction coordinate (RC) optimisation, the input features were preprocessed using tICA; in all single-fibre (tetra-nucleosome) cases, we retained 100 tICA eigenvectors (dim = 100), which served as input for reaction coordinate training at a lag time *τ* ^FMRC^ = 250 ns. The rest of training parameters followed the standard FMRC algorithm training protocol [38] (https://github.com/Mingyuan00/Flow_Matching_for_RC). We then clustered simulation frames into 1500 microstates in this optimal RC and constructed an MSM with a lag time of 250 ns. For the tetra-nucleosome condensate system, we considered both chromatin arrays in the condensed phase (*N* = 54) and all arrays in the simulation box (*N* = 81), using the same feature description as in the single-fibre analysis. For these condensate systems, 100 tICA eigenvectors were retained and used for FMRC training at a lag time of 25 ns. The resulting ensembles were then clustered into 500 microstates, and MSMs were constructed in the optimised RC space using a lag time of 250 ns. The MSM was constructed using a detailed-balance-constrained maximum-likelihood (ML) estimator, implemented as `MaximumLikelihoodMSM` in the Deeptime library [62]. All subsequent validation analyses were also performed using Deeptime. RC learning was carried out in PyTorch (version 2.7.1 with CUDA 12.6 support). The parameters used for all systems in this work are summarized in the Supplementary Information.

### Analyses

#### MSM reweighting and feature correlation

The MSM-reweighted free-energy surface (FES) was represented in the physically interpretable coordinate space (*d*_13_, *d*_24_) by assigning each trajectory frame a statistical weight derived from the MSM equilibrium distribution. After constructing and validating the MSM transition matrix **T**(*τ*) at lag time *τ*, the stationary distribution ***π*** was obtained as the left eigenvector associated with *λ*_1_ = 1: ***π***^T^ = ***π***^T^ **T**(*τ*). This vector provides the equilibrium probabilities of the microstates. For each trajectory frame *k*, let *m*_*k*_ denote its assigned microstate. The frame weight was then defined as

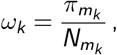

where 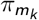 is the stationary probability of the microstate containing frame *k*, and 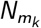 is the total number of frames assigned to that microstate. The (*d*_13_, *d*_24_) plane was discretized into bins indexed by (*x*, *y*). The reweighted probability in each bin was computed by summing frame weights, By accumulating the weighted counts, we obtained the probability distribution

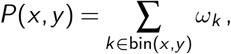

and the corresponding free energy was evaluated as

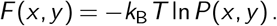

Input features were related to the slow dynamical modes of the MSM by computing ***π***-weighted Pearson correlations between microstate-averaged feature values and the second and third right eigenvectors of the MSM transition matrix. Correlations were computed using the SciPy library [63]. For each feature *l*, its mean value in microstate *m* was first computed as:

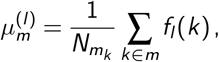

where 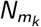 and ***π*** follow the definitions above, and correlations were weighted by the stationary distribution ***π***.

#### Reference structure preparation

The reference structures required by the RMSD function in PLUMED were generated by an in-house clustering Python script. Specifically, for each four-nucleosome structure, we computed six pairwise inter-nucleosomal distances {*d*_12_, *d*_13_, *d*_14_, *d*_23_, *d*_24_, *d*_34_} using centre-of-geometry positions for each nucleosome. These were transformed into symmetrized features invariant under the label permutation (1, 2, 3, 4) ↔ (4, 3, 2, 1), following the approach of Ding and Zhang [8]. K-means clustering was then performed on the six-dimensional symmetrized distance features using the `deeptime.clustering` module. Three clusters were obtained, and reference structures were extracted from the most populated cluster.

#### Nucleosome-nucleosome contact analysis

Nucleosome–nucleosome contacts were identified following a protocol adapted from our previous work [11] and analysed using MDtraj (1.10.3) [64]. For each nucleosome, we computed the nucleosome-core centre-of-mass (CoM) and the corresponding superhelical axis. Two nucleosomes (*r*_*i*_, *r*_*j*_) were defined as interacting when their inter-CoM distance *d*_*ij*_ = |*r*_*i*_ − *r*_*j*_| < 13 nm. To classify the orientation of each interacting pair, we considered the unit vector 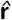 connecting the two CoMs and the unit superhelical axes 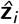 and 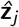 of nucleosomes *i* and *j*. Three angular quantities were computed:

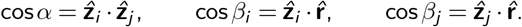

Contacts were classified as follows:

- **Face–face:** (*α* < 45^*◦*^ or *α* > 135^*◦*^), and at least one nucleosome axis is approximately aligned with the inter-CoM vector (*β*_*i*_ or *β*_*j*_ outside [45^*◦*^, 135^*◦*^]), or if *d*_*ij*_ < 11 nm.
- **Side–side:** (*α* < 45^*◦*^ or *α* > 135^*◦*^), but neither nucleosome axis is aligned with the inter-CoM vector (*β*_*i*_, *β*_*j*_ ∈ [45^*◦*^, 135^*◦*^]) and *d*_*ij*_ ≥ 11 nm.
- **Face–side:** all remaining contacts.

#### Density profile analysis

The nucleosome density profile was computed along the x-axis, the assumed primary axis of phase separation. For each frame, nucleosome positions were determined from their CoMs, and the overall system centre was subtracted to remove translational drift. The resulting x-coordinates were wrapped into [−250, 250) nm and discretized into 12.5 nm intervals along the 500 nm periodic box. Per-frame histograms of nucleosome counts were then averaged to obtain the mean density profile, with the standard deviation across frames used to quantify fluctuations. Bins with a mean count above 36 were classified as condensed. A tetra-nucleosome was retained as condensed if at least one of its four nucleosomes occupied a condensed bin in ≥75% of simulation frames. Using this criterion, 54 of the 81 tetra-nucleosome arrays in the system were identified as belonging to the condensed phase (Figure S11).

## DATA VISUALISATION AND AVAILABILITY

Structures were visualized using the Open Visualization Tool (OVITO Pro) [65] and VMD [66]. Data were plotted using Matplotlib (version 3.9.4) and NetworkX (version 3.2.1) [67]. Simulation data are available from the corresponding author upon reasonable request.

## CODE AVAILABILITY

All Python scripts used in this work are available in the GitHub repository (https://github.com/yifangchen7/chromatin-fiber-msm). The chemically-specific coarse-grained simulation package OpenCGChromatin is available at https://github.com/CollepardoLab/OpenCGChromatin. The minimal coarse-grained model is available at https://github.com/CollepardoLab/CollepardoLab_Chromatin_Model.

## ACKNOWLEDGEMENTS

Research in the Collepardo-Guevara lab is supported by the UK Research Innovation (UKRI) Engineering and Physical Sciences Research Council (EPSRC) [EP/Z002028/1], following funding from the European Research Council (ERC) Consolidator Grant “ChromatinDroplets” under the European Union’s Horizon Europe research and innovation programme. We acknowledge EuroHPC Joint Undertaking for awarding access to MareNostrum5 at Barcelona Supercomputing Center (BSC), Spain [EHPC-REG-2025R01-166]. Y.C. would like to acknowledge the Oliver Gatty Studentship Fund for doctoral funding.

## Notes

### Competing Interest Statement

R.C.G. and J.R.E. are co-founders of Phasica Biosciences S.L. All other authors declare no competing interests

